# Parallel Molecular Data Storage by Printing Epigenetic Bits on DNA

**DOI:** 10.1101/2023.12.15.571646

**Authors:** Cheng Zhang, Ranfeng Wu, Fajia Sun, Yisheng Lin, Yizheng Zhang, Yuan Liang, Yiming Chen, Jiongjiong Teng, Zichen Song, Na Liu, Qi Ouyang, Long Qian, Hao Yan

## Abstract

DNA storage has shown potential to transcend current silicon-based data storage technologies in storage density, lifetime and energy consumption. However, writing large-scale data directly into DNA sequences by *de novo* synthesis remains uneconomical in time and cost. Inspired by the natural genomic modifications, in this work, we report an alternative, parallel strategy that enables the writing of arbitrary data on DNA using premade nucleic acids. With epigenetic modifications as information bits, our system employed DNA self-assembly guided enzymatic methylation to perform movable-type printing on universal DNA templates. By programming with a finite set of 700 DNA movable types and 5 templates, we achieved the synthesis-free writing of ∼270000 bits on an automated platform with 350 bits written per reaction. The data encoded in complex epigenetic patterns were retrieved high-throughput by nanopore sequencing, and algorithms were developed to finely resolve 240 modification patterns per sequencing reaction. Our framework presents a new modality of DNA-based data storage that is parallel, programmable, stable and scalable. Such a modality opens up avenues towards practical data storage and dual-mode data functions in biomolecular systems.

## Introduction

The dramatically expanding global data-sphere has posed an imminent challenge on large-scale data storage and an urgent need for better storage materials^1,2^. Inspired by the way genetic information is preserved in nature, DNA has been recently considered a promising biomaterial for digital data storage owing to its extraordinary storage density and durability^3-8^. In current DNA storage, data is typically transcoded into a sequence of nucleotide bases, and writing largely depends on *de novo* chemical synthesis in which nucleotide bases are added one at a time in predetermined orders^9-13^. Although technological upgrades of *de novo* DNA synthesis have been driving a continuous increase in throughput and efficiency^14-18^, the serial synthesis of individual DNA molecules essentially limits the writing speed and the length of synthesized DNA, and prevents a substantial cost reduction in data writing^18-22^.

In this work, we develop a completely different framework for data writing and use it to establish the largest scale of storage so far (∼270000 bits) based on unconventional molecular information bits. We propose a parallel DNA storage strategy based on DNA self-assembly guided enzymatic methylation to selectively write epigenetic bits (epi-bits), much as typography to press information on papers (Fig. 1B)^29^. First, a universal ssDNA carrier and a library of complementary short ssDNA bricks are predesigned for programmable typesetting (Fig. 1C). Then, arbitrary epi-bit information can be typeset by assembling the premade library onto the identical loading sequence of the DNA carriers. Next, the modification information is parallelly and stably “printed” on the DNA carriers through selective methylation by the methyltransferase DNMT1 (Fig. 1D). Finally, numerous epi-bit streams stored on different carrier molecules is retrieved high-throughput by one-pot nanopore sequencing (Fig. 1E). The strategy enables the parallel writing of arbitrary data in DNA using premade nucleic acids instead of *de novo* synthesis. This enzymatic printing process potentially reduces the cost and time beyond the limit of chemical synthesis, and the fidelity of data writing is endowed by highly specific brick-template DNA assembly.

**Figure 1.**
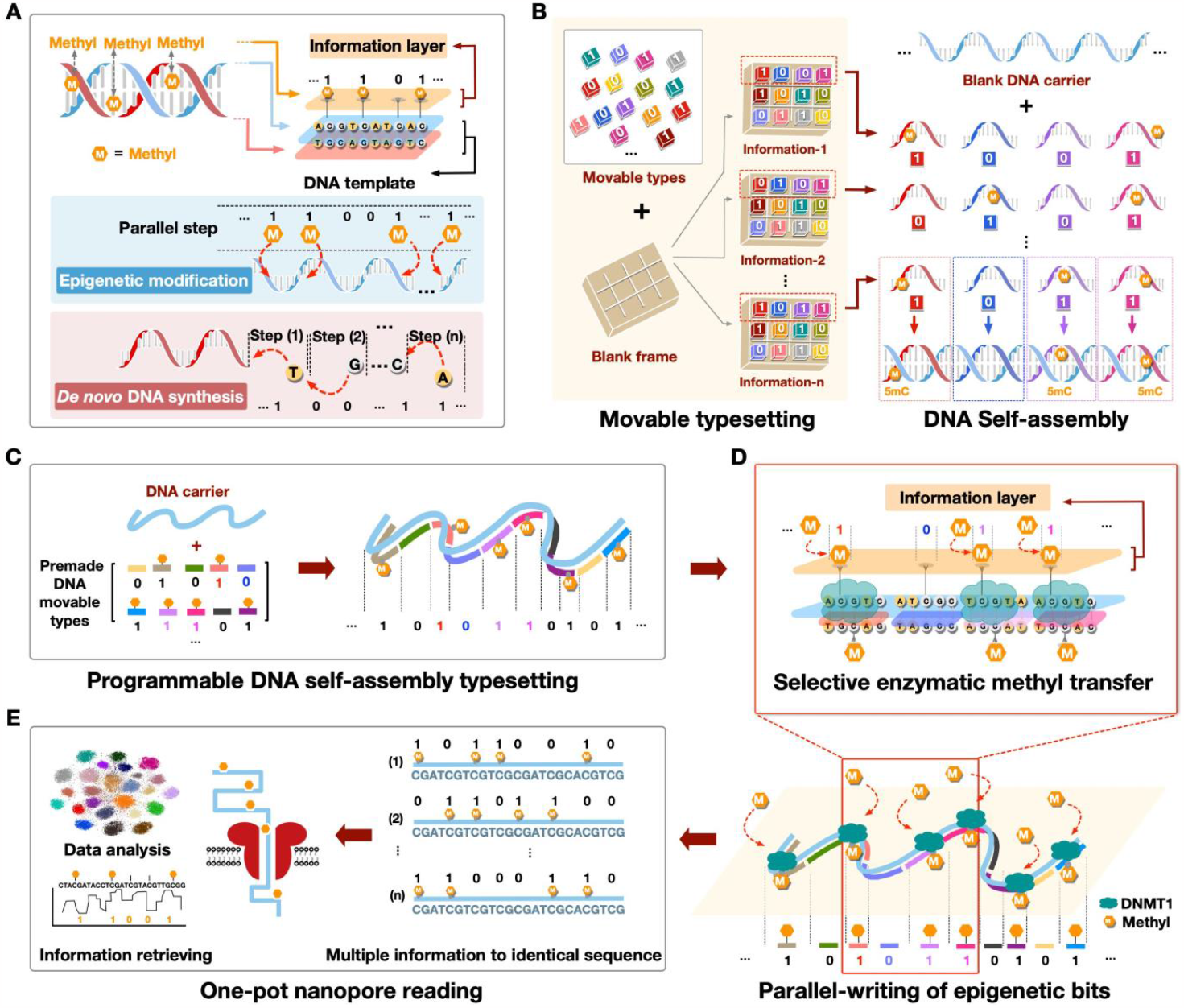
Schematics of the epi-bit DNA storage. (A) Illustrations of the epigenetic information storage. The modification information is typeset by specifically programing DNA movable types carrying epi-bits (B&C) and written by selective enzymatic methyl transfer (D). (E) Nanopore sequencing of modified templates and methylation calling.

## Results

### *In vitro* selective epi-bit writing and reading

To test *in vitro* epi-bit writing, we first implemented selective transfer of a single-bit methyl group onto the DNA template. In the design, the methylation site, a CpG dinucleotide, is set in the middle of an ssDNA brick p or q with a methylated or unmethylated cytosine, respectively (Fig. 2A). DNMT1 specifically recognizes the hemimethylated site within DNA p and transfers a methyl group from S-adenosylmethionine (SAM) to the opposite cytosine in the DNA template^27^, achieving the writing of epi-bit value 1. In contrast, for an unmethylated brick q, no methyl group transfer happens and the DNA template remains in the epi-bit 0 state (Fig. 2A). To test the efficiency of such selective epi-bit writing, we developed a molecular beacon (Fig. 2B). The template DNA b1 and the guide DNA a1 were initially hybridized to bring a quencher and a fluorophore in proximity (fluorescent OFF state). Only when the DNA complex a1b1 contained a quadruply methylated GCGC site, can it be cleaved by the restriction endonuclease GlaI, leading to the fluorophore release and an increase in fluorescence^30^. We started with singly methylated DNA template b1 (G5mCGC) and tested whether the methyl group can be transferred by DNMT1 (NEB®) at the second cytosine when guided by DNA a1 (G5mCG5mC, epi-bit “1”) and a2 (GCG5mC, epi-bit “0”), respectively (Fig. 2B). From the results of gel and fluorescence assays, it is clear that only the DNA a1 induced methyl group transfer, indicating the precise selective writing of a single epi-bit (Fig. 2C&D).

**Figure 2.**
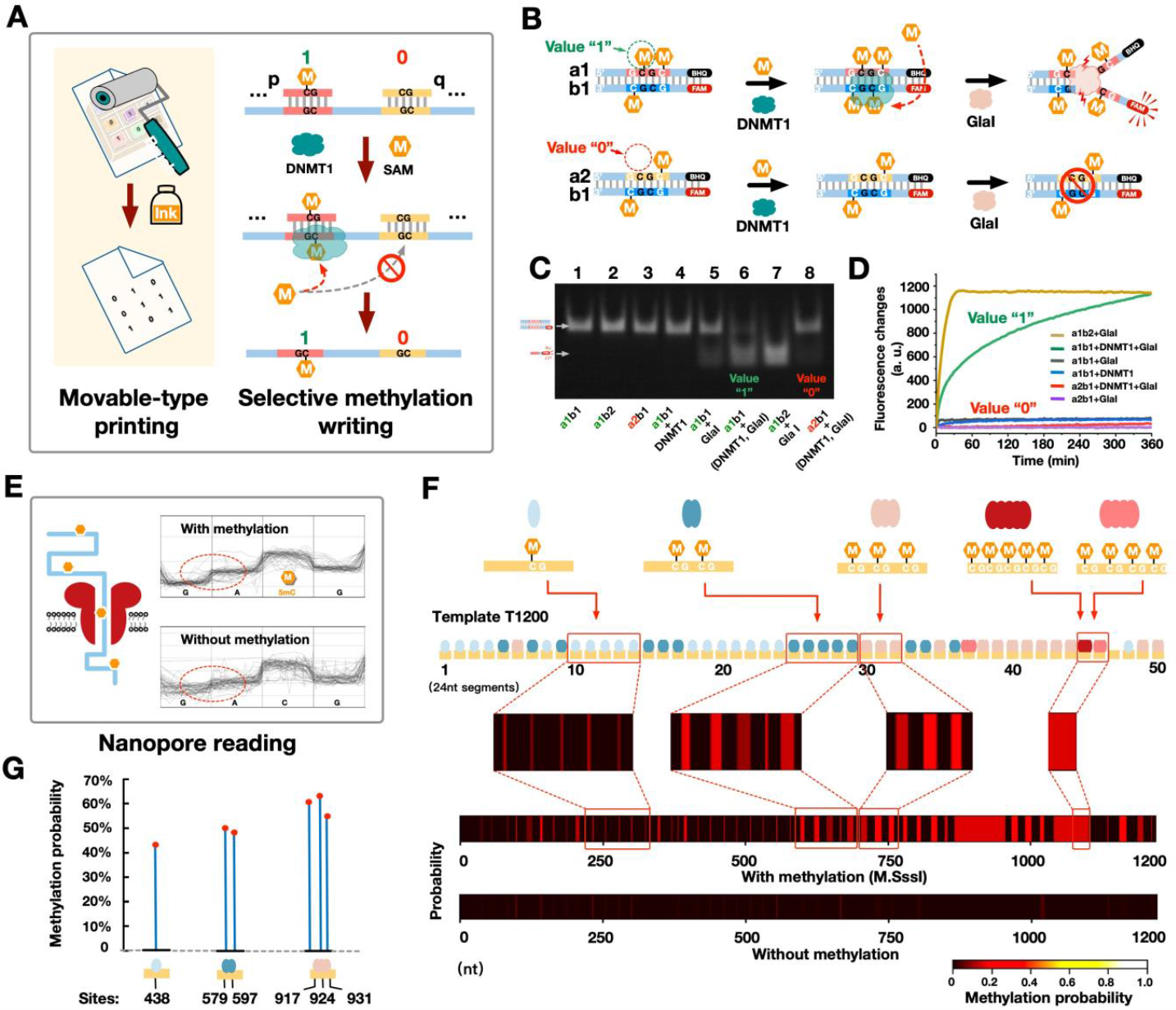
Test on the epi-bit writing and the nanopore reading. (A&B) The illustrations of the principle and the design of single-bit methylation writing assay. (C) Gel shift assay detecting different methylation states. (DNA b2: positive control of double methylated DNA b1). (D) Fluorescent detection of different methylation states. (E) Illustration of the epi-bit nanopore reading. (F&G) Methylation calling of the nanopore signals of templates treated by M.SssI (F) and DNMT1 (G) using Nanopolish and Megalodon, respectively.

For epi-bit reading, we explored the nanopore sequencing technology (Fig. 2E). DNA template T1200 (1200 nt) was designed with 101 CpG dinucleotides as potential methylation sites and treated by a highly efficient 5mC methyltransferase M.SssI ^31^(NEB®, Fig. 2F). The CpG dinucleotides were designed to distribute into segments along the template T1200, yielding five kinds of segments I, II, III, IV and V containing 1, 2, 3, 4 and 5 CpG sites, respectively (Fig. 2F). The nanopore current signals of M.SssI treated, CpG fully-methylated DNA templates exhibited significant signal shifts around methylation sites (Fig. 2E). Methylation calling by Nanopolish reliably detected all segment types even when the segment contained only one 5mC ^32^ (Fig. 2F). In contrast, almost no significant methylation was called from the templates untreated with M.SssI. Additionally, selective epi-bit writing was implemented with DNMT1 and ssDNA bricks. The nanopore signals called by Megalodon ^33^ exhibited remarkable resolution in distinguishing three closely spaced epi-bits in a brick segment (Fig. 2G).

### Programmable epi-bit typesetting through DNA self-assembly

To program arbitrary epi-bit data, a DNA self-assembly based typesetting strategy was developed (Fig. 3A(i)). A typical DNA movable type was designed as a ssDNA brick with one CpG dinucleotide in the middle as the epi-bit location. The epi-bit value (1 or 0) of movable types is governed by the presence or absence of a C5-methylated cytosine in the CpG dinucleotide (Fig. 3A(ii)). By selecting specific combinations of DNA movable types to assemble with ssDNA carriers, the epi-bits on movable types were parallelly aligned to establish a methylation pattern representing the coded information. Here, DNA carrier T960 (960 nt) was established to store 32 bits information in its loading sequence with a total of 72 kinds of movable types. We tested the operation by assembling carrier T960 with 10 to 36 movable-types. In gel results, DNA carriers shifted to low mobility bands stepwise with increasing numbers of movable types, indicating successful DNA typesetting (Fig. 3B).

**Figure 3.**
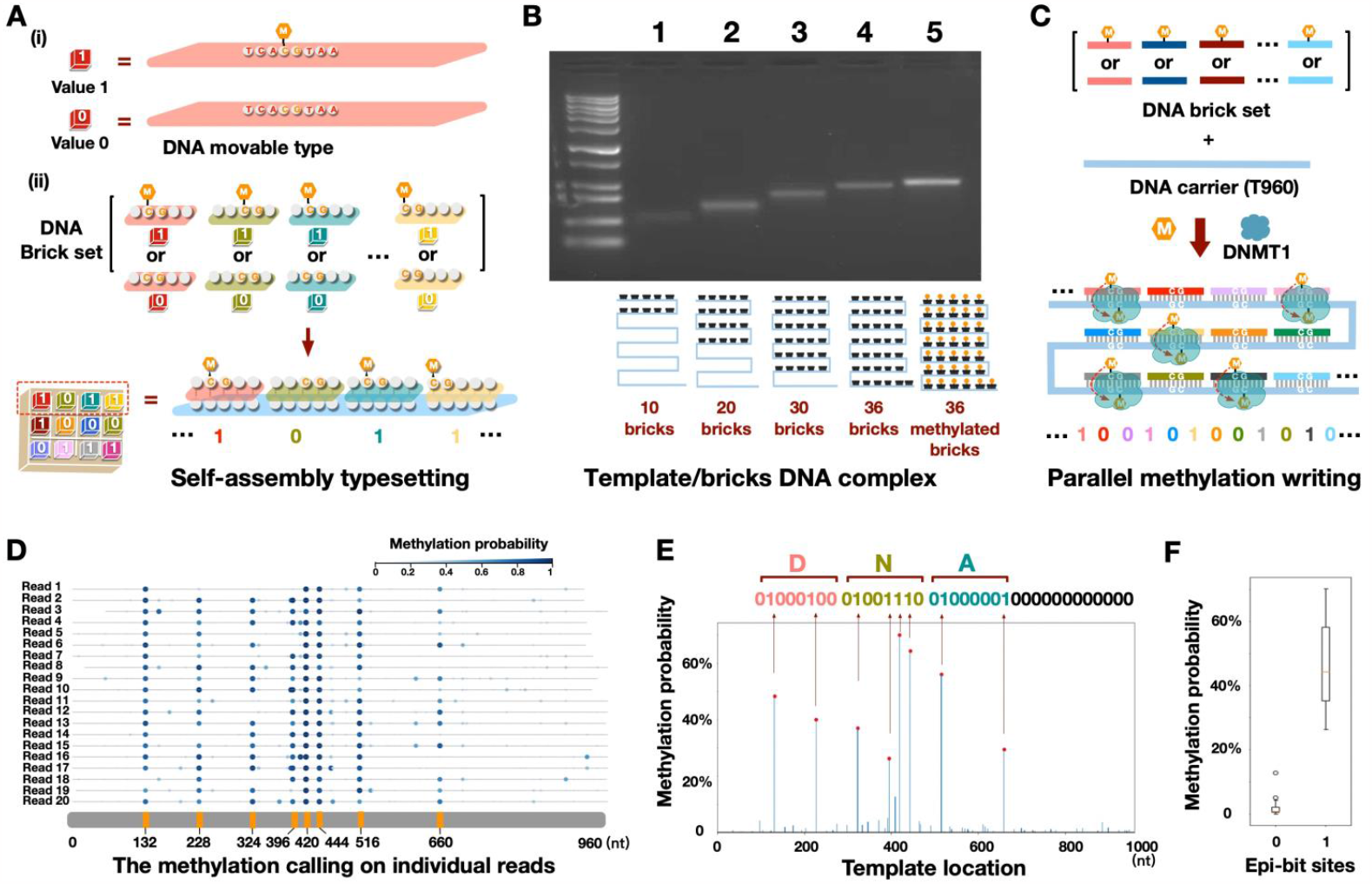
Programmable DNA typesetting and parallel epi-bit writing. (A) Diagram of programmable self-assembly typesetting. (B) The gel results of the 960 nt DNA carriers assembled with varying numbers of DNA bricks. (C) Diagram of parallel enzymatic epi-bit writing. (D) The predicted methylation probabilities on twenty sequencing reads of DNA carrier encoding “DNA”. (E) The overall nanopore sequencing retrieved epi-bit information of “DNA”. (F) The predicted methylation probability distribution at epi-bit 0 vs. epi-bit 1 sites.

Next, parallel epi-bit writing was implemented by using DNMT1 to transfer methylation information from the movable types to the DNA carrier (Fig. 3C). As a demonstration, we wrote the ASCII code for letters “DNA” (24 bits) to one T960 DNA carrier. After nanopore sequencing and methylation calling, the information was retrieved with negligible background noise (Fig. 3 D&E). As signal distributions were distinct for epi-bit 1 sites *versus* epi-bit 0 sites (Fig. 3F), it was possible to apply a threshold for accurate information retrieval.

### Enlarged epi-bit storage capacity by DNA sequence-based barcoding

Because nanopore sequencing resolves both sequence and modification information, we first scaled up the storage experiment by writing 800-bit information onto 25 kinds of barcoded DNA carriers with a universal DNA template sequence using the same set of 72 movable types (Fig. 4 A to C). Structure drawings of four DNA bases and four of their modified derivatives were first transformed by bitmap coding and sparsified to reduce the number of epi-bit 1s (Fig. 4A). The data were then allocated to different DNA carriers. The DNA carriers were designed to each have two functional regions: 1) a unique barcode region for indexing and 2) a universal 960 nt template as the loading sequence for epi-bit writing (Fig. 4B). The two regions were individually generated, aligned by an ssDNA linker and ligated (Fig. 4C). For writing, all DNA carriers were treated by DNMT1 as guided by specific combinations of DNA movable types representing the allocated data, which totaled 800 bits (Fig. 4C&D).

**Figure 4.**
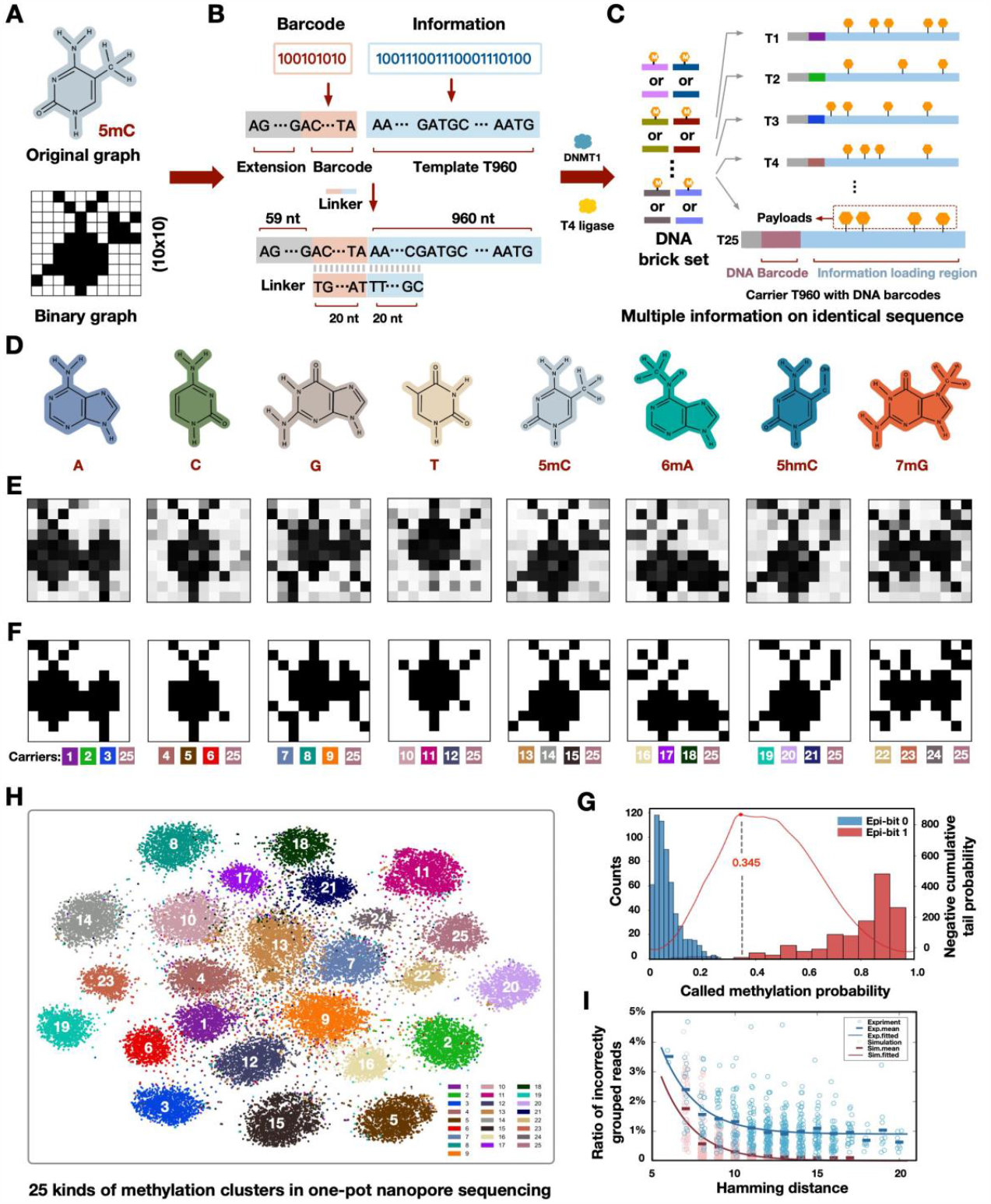
Enlarged epi-bit data storage and read by one-pot sequencing. (A) Processing of the original image data. (B) Preparation of barcoded DNA carriers. (C) Writing of multiple epi-bit information. (D) Stored images of DNA bases and derivatives. (E) Images recovered by called methylation probabilities in greyscale. (F) Binarized image recovery based on the threshold of methylation calling. (G) Determination of the threshold for methylation calling. (H) K-means clustering of all sequencing reads based on called epi-bit patterns. (I) Total barcode mixing fractions as a function of the hamming distance between pairs of epi-bit patterns.

For data retrieval, one-pot nanopore sequencing of mixed DNA carriers yielded read signals which were first grouped by the methylation patterns, screened for the correct barcodes, and then called for methylation collectively. Fig. 4E&F show the recovery of all original images from sequenced epi-bit streams. Comparison of the methylation probability distribution at each epi-bit site revealed differential discriminability between 0 and 1, likely attributed to local context effects on methylation efficiency, variant self-assembly efficiencies owing to the DNA sequence design, or contextual signal detection in nanopore reads^34^. From these distributions, a methylation calling threshold was obtained as 0.345 which resulted in an error rate of 0.625% (Fig. 4G).

Besides errors in methylation calling, errors could have originated from the promiscuous ligation between barcodes and T960 loading templates. To analyze relative contributions of the two error sources, K-means clustering (*k* = 25) of reads purely based on their methylation patterns was further analyzed. In Fig. 4H, each cluster corresponds to one segment of data written on a specific barcoded DNA carrier. Nevertheless, an average 25.5% fraction of barcodes within the clusters were incorrectly identified. We numerically simulated barcode mixing due to methylation miscalling as a function of epi-bit distances at the experimentally measured per read miscalling rate E_meth_ = 0.3 per site. After removing the effect of miscalling from the total misidentified barcode fractions, the promiscuous ligation rate was estimated to be E_lig_ = 0.9% regardless of epi-bit distances (Fig. 4I). Nonetheless, without the reference of methylation patterns, information retrieval based solely on barcodes had a mere error rate of 1.75%.

### Large-scale storage with high bit parallelism

To leverage the parallel nature of the epi-bit writing mechanism, we implemented large-scale data storage with multiple DNA templates and denser epi-bit sites (Fig. 5A). We define the bit throughput or parallelism, an essential feature of data writing, as the number of bits written in a single minimal reaction system per data-writing cycle. For example, *de novo* synthesis has a bit throughput of ∼1, and for the above experiments the bit throughput was 32. In the next set of large-scale experiments, we stored the image of a tiger rubbing (16833 bits) from the Han dynasty in ancient China and the colored picture of a panda (252504 bits) (Fig. 5A (i)). Here, 5 kinds of 1300 nt DNA templates (L1 to L5) selectively hybridized with 175 DNA bricks (from a movable-type set of 700 bricks), achieving 350-bit parallelism per writing reaction (Fig. 5A (ii)). Notably, reactions were labeled by epi-bit barcodes stored in the modification layer instead of ligated DNA barcodes in order to reduce complexity and streamline experimental protocols (Fig. 5A (iii)).

**Figure 5.**
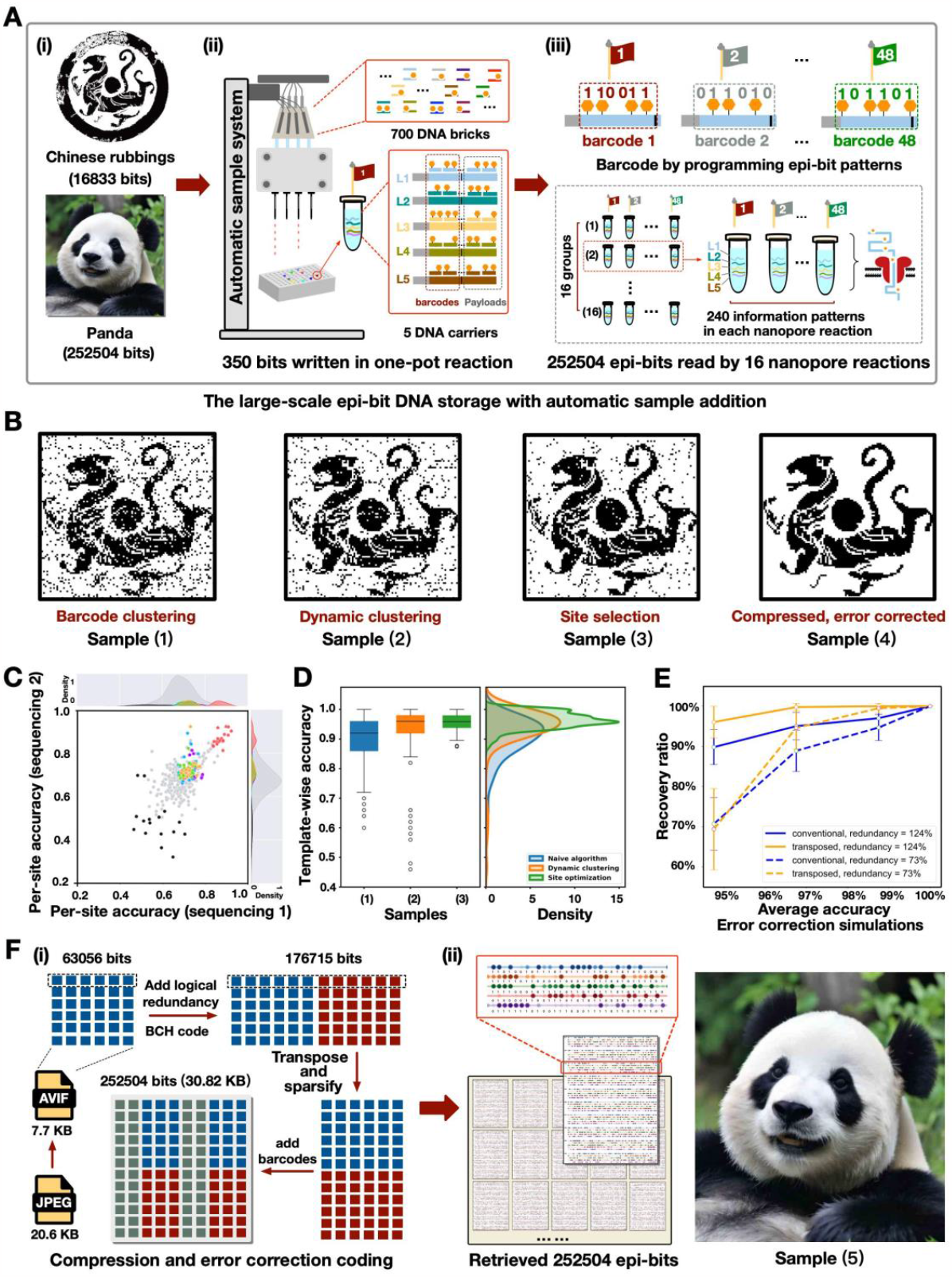
Large-scale storage with high bit parallelism by epi-bit barcodes. (A) The process of large-scale epi-bit DNA storage. (B) The recovered rubbings images of samples (1) to (4) with step-wise improved writing-reading pipelines. (C) The epi-bit per-site accuracy between two independent sequencing samples of the original tiger encoding. (D) The distribution of template-wise accuracy of samples (1) to (3). (E) Simulations of the error correction capabilities of different algorithms for the storage of the panda image. (F) The compression and error correction coding method to store the panda image.

We firstly stored the image of a tiger rubbing by bitmap encoding (16833 bits distributed to 48 barcodes, Fig. 5B, sample (1)). After nanopore reading, methylation calling on grouped nanopore reads based on barcode information (i.e., the barcode clustering method) yielded an accuracy of 90.14% primarily due to the noisy single-read methylation probabilities (E_meth_ = 0.3) at barcode sites (Fig. 5B, sample (1)). Therefore, we sought to improve read classification by an iterative dynamic clustering algorithm that also consulted data at information sites, which increased the accuracy to 93.65% (Fig. 5B, sample (2)). Next, as we observed significant but consistent variations in per-site methylation discriminability, site selection was implemented, in which sites with low accuracies (2.05% of all sites) were discarded from coding and sites ranked top in accuracy in each template served barcode sites for a third storage experiment (Fig. 5C). Site selection significantly improved the data retrieval accuracy to an average of 96.33% with much narrower distributions among templates (Fig. 5B, sample (3) and Fig. 5D). Finally, we developed a compression and error correction coding scheme to achieve full data recovery from one-pot nanopore sequencing reads (Fig. 5B, sample (4)). *In silico* evaluation of various error correction strategies suggested transposition effectively dispersed clustered errors to recover the original information from up to 5.3% error rate (Fig. 5E).

Lastly, we integrated the above strategies to store the image of a panda, which totaled 252504 bits (15-fold of the tiger image) after compression, error corrective encoding and barcoding (Fig. 5F(i)). The data were written in 756 reaction wells by the automatic platform. For retrieval, every 48 wells (240 epi-bit patterns stored on DNA templates) were sequenced in one batch and the dynamic algorithm yielded an average accuracy of 97.47%. With error corrective decoding, the image was perfectly restored (Fig. 5F(ii)).

## Discussion

In this work, we reported an efficient and scalable movable-type DNA storage framework that allows arbitrary epi-bit information to be stably written onto DNA templates with a premade set of DNA movable types and the methyltransferase DNMT1. To our knowledge, this work showcases the largest scale of storage among reported unconventional DNA data storage strategies. We demonstrated our framework by the parallel writing of ∼270000 bits data on 5 sets of DNA carriers. The data stored on a mixture of DNA carriers were retrieved by nanopore sequencing at accuracies of ∼99.4% and ∼97.47%.

Overall, this work demonstrates potential directions in parallel molecular information processing with prefabricated modularity. Combining DNA self-assembly assisted programming and a myriad of enzymatic modifications, it is possible to realize diverse DNA storage and computation functions for practical and functionalized molecular data systems.

